# The apple palmitoyltransferase MdPAT16 regulates sugar content via an MdCBL1-MdCIPK13-MdSUT2.2 pathway

**DOI:** 10.1101/2020.05.28.121970

**Authors:** Han Jiang, Qi-Jun Ma, Ming-Shuang Zhong, Huai-Na Gao, Yuan-Yuan Li, Yu-Jin Hao

## Abstract

Protein palmitoylation, a post-translational protein modification, plays an important role in the regulation of substrate protein stability, protein interactions, and protein localization. It is generally believed that there are two mechanisms of palmitoylation: one by acyl-CoA and the other by protein acyltransferase (PAT). In this study, an MdPAT family member, MdPAT16, was identified and shown to have palmitoyltransferase activity. We found that this gene responded to salt stress and that its expression improved plant salt resistance. MdPAT16 was shown to interact with MdCBL1 and stabilize MdCBL1 protein levels through palmitoylation. MdPAT16 further regulated apple sugar content by stabilizing the MdCIPK13-MdSUT2.2 protein complex. We found that the N-terminal sequence of MdCBL1 contains a palmitoylation site and that the N-terminal deletion of MdCBL1 leads to changes in protein stability and subcellular localization. Finally, exogenous salt stress increased the interaction of MdPAT16 with MdCBL1 and the sugar content in apple. These findings suggest that MdPAT16 functions as a stable means for the palmitoylation of downstream protein. It may be a missing link in the plant salt stress response pathway and have an important impact on fruit quality.

## INTRODUCTION

Sugars play multiple important roles in diverse biological processes, such as physiological metabolism, growth, and developmental stage transitions. Sugars not only supply energy for plant growth and development, but also act as important determinants of fruit quality and commodity value. The content, type, and ratio of sugars directly or indirectly determine fruit flavor, color, and other quality traits. Sugars also function as signaling molecules to regulate flowering time, nitrogen metabolism, anthocyanin accumulation, and other processes in various plant species (Ohto et al., 2001; Jonassen et al., 2008; Sun et al., 2019; Hu et al., 2016; Liu et al., 2019).

Sugars are also used for osmotic adjustment in response to various abiotic stresses. Salinity, drought, and low temperature usually result in the accumulation of sugars (Krasensky et al., 2012). Under long-term salt stress, plants enhance their salt tolerance in response to dehydration and osmotic stress (Chaves et al., 2009). Early salt stress is similar to drought and is mainly affected by ion toxicity and osmotic stress. Therefore, increasing plant tolerance to the early stages of salt stress mainly involves preventing salt from entering the cytoplasm and lowering cell water potential through osmotic adjustment(Sultana et al., 1999; Parida and Das, 2005; Munns and Tester, 2008). The sugars produced by photosynthesis provide necessary energy for the growth and development of tissues in response to environmental stimulation (Lv et al., 2008; Yao et al., 2010). When sugars accumulate to a relatively high level in the vacuole, this produces a high bloating pressure (Gibson, 2005; Moustakas et al., 2011; Rasheed et al., 2011). Therefore, sugars also affect osmotic potential and participate in the response to water and salt stresses.

In *Arabidopsis*, sugars participate in abiotic stress responses by affecting osmotic potential, and increased contents of soluble sugar, anthocyanins, and proline occur under stress (Moustakas et al., 2011). In wheat, the contents of glucose, fructose, sucrose, and fructan increase significantly under drought or salt stress (Kerepesi et al., 2000). In apple, the CBL-interacting protein kinase (CIPK) MdCIPK22 interacts with and phosphorylates the sucrose transporter MdSUT2.2 to mediate drought tolerance and sugar accumulation (Ma et al., 2019a). The ABA-related transcription factor MdAREB2 directly binds to the promoter of MdSUT2.2, which plays an important role in ABA-induced sugar accumulation (Ma et al., 2017a). Meanwhile, MdCIPK22 also interacts with and phosphorylates MdAREB2, thereby promoting its transcription (Ma et al., 2017b). Therefore, MdCIPK22 both directly and indirectly regulates the downstream protein MdSUT2.2 and sugar accumulation in apple. Another CIPK family gene, MdCIPK13, may also regulate sugar content and salt tolerance (Ma et al., 2019b). However, the upstream regulatory pathways for MdCIPK13 in response to salt stress remain unclear.

S-acylation, also called S-palmitoylation, involves the binding of a 16-carbon palmityl group to a specific protein cysteine residue through a thioester bond. S-palmitoylation regulates dynamic membrane localization, stability, and transport of proteins between different cellular compartments. It also regulates protein function and protein-protein interactions. Although the mechanism of palmitoylation remains unclear in plants (Hemsley et al., 2013), it is well known that palmitoylation occurs in different membrane structures, such as the endoplasmic reticulum (Batistič et al., 2008), Golgi (Zeng et al., 2007), the plasma membrane (Sorek et al., 2007), and the tonoplast (Batistič, et al., 2012; Zhou et al., 2013). Palmitoylation of proteins also occurs spontaneously without enzyme catalysis. However, in this process, a large number of reaction substrates must be provided (Bizzozero et al., 1987; Duncan and Gilman, 1996). As organisms fail to provide such large amounts of substrate, spontaneous protein palmitoylation is unlikely to occur *in vivo*. The common understanding is that protein palmitoylation is catalyzed by a series of enzymes called protein S-acyl transferases (PATs) (Batistič, et al., 2012; Zhou et al., 2013).

Palmitoyl transferase was initially discovered in *Saccharomyces cerevisiae*, and it has been extensively studied in mammals and yeast (Sun et al., 2004; LaGrassa and Ungermann, 2005). The first PAT function genes to be characterized were the Erf2-Erf4 complex and Ark1, which promote palmitoylation of yeast Ras2 protein and Yck2 (yeast casein kinase 2) kinase, respectively (Feng and Davis, 2000; Qi et al., 2014). Both Erf2 and Ark1 have a conserved DHHC (Asp-His-His-Cys) motif in the cysteine residue aggregation domain (CRD), as well as zinc finger structural features (González-Siso et al., 2009; Hou et al., 2005; Ohno et al., 2006; Subramanian et al., 2006). Generally, proteins rich in DHHC motifs have PAT activity. These motifs are important not only for PAT activity, but also for palmitoylation of the DHHC protein itself (Qi et al., 2014).

In plants, the first reported palmitoyl transferase was TIP1 (TIP GROWTH DEFECTIVE 1) in Arabidopsis. It is a member of the palmitoyl transferase family with an ankyrin repeat sequence and is generally expressed in roots, leaves, inflorescence stems, and flowers. TIP1 regulates protein hydrophobicity and affects protein-membrane binding, signal transduction, and intracellular vesicle trafficking (Hemsley et al., 2005). In addition, another Arabidopsis palmitoyltransferase, *AtPAT10*, has palmitoyltransferase activity and participates in the regulation of cell expansion and cell division. It also enhances reproductive capacity (Qi et al., 2014). The AtPAT10 protein localizes to the Golgi apparatus and the tonoplast (Qi et al., 2013; Zhou et al., 2013). The phenotypes of three *AtPAT10* T-DNA insertion mutants are consistent and include defects in cell expansion and cell division, as well as hypersensitivity to salt stress. *AtCBL2* and *AtCBL3* have been identified as potential substrates of *AtPAT10* and shown to regulate salt tolerance (Zhou et al.,2013). However, it remains unclear how PATs regulate sugar accumulation in response to salt stress.

In this study, a palmitoyltransferase family member, *MdPAT16*, was identified in apple. Functional complementation and S-acylation experiments demonstrated that MdPAT16 has palmitoyltransferase activity, and subsequent experiments characterized its functions in sugar accumulation and salt stress tolerance. Its interacting protein MdCBL1 was also identified and characterized. Finally, the response of MdPAT16 to salt stress and its interactions with MdCBL1 to regulate sugar accumulation and modulate salt tolerance were characterized.

## RESULTS

### MdPAT16 promotes sugar accumulation and enhances salt tolerance

Esculin staining of roots demonstrated that an appropriate concentration of NaCl clearly promoted sucrose transferase activity in apple roots, thereby increasing plant sugar content (Fig. S1). RNA-seq analysis further demonstrated that numerous genes associated with sugar biosynthesis and transportation were markedly upregulated. In addition to sugar-associated genes, the PAT family member *MdPAT16* was also transcriptionally upregulated in response to NaCl treatment (Fig. 1).

**Figure 1.**
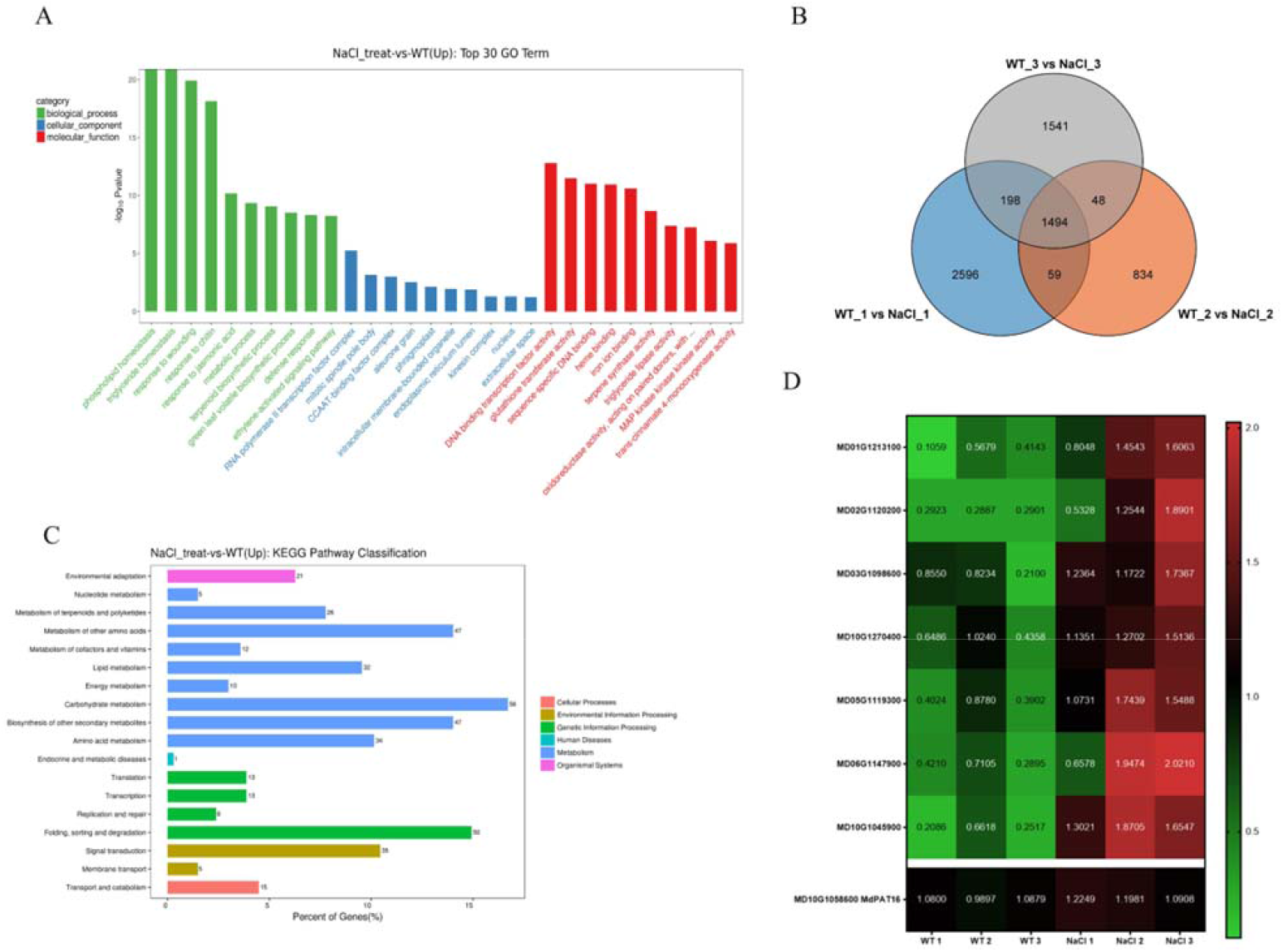
RNAseq analysis of GL-3 tissue culture seedlings with and without NaCl treatment. **(A)** Gene ontology enrichment of upregulated genes in NaCl-treated and WT seedlings, presented according to −log_10_P-value. **(B)** Venn diagram of the genes upregulated in NaCl-treated and WT seedlings (log_2_FoldChange > 2). **(C)** KEGG pathway classification of upregulated genes. **(D)** Normalized heatmap of sugar-related genes, including amino sugar and nucleotide sugar metabolism(MD01G1213100,MD02G1120200), starch and sucrose metabolism(MD03G1098600, MD10G1270400), sucrose transmembrane transporter(MD10G1045900), salt stress response (MD05G1119300, MD06G1147900), and MdPAT16(MD10G1058600).

To further characterize the function of *MdPAT16* in response to salt stress, a pMdPAT16-GUS vector was transiently transformed into apple shoot cultures. The transgenic shoot cultures were then treated with NaCl or H_2_O. After GUS staining, NaCl-treated shoot cultures exhibited higher GUS activity than water-treated controls (Fig. S2). To further verify that MdPAT16 functions in salt-induced sugar accumulation, *Agrobacterium rhizogenes*-mediated genetic transformation was performed to obtain MdPAT16 overexpression and suppression transgenic roots. Two weeks after NaCl treatment, MdPAT16-OVX roots grew much better and MdPAT16-Anti roots grew poorly compared with the empty vector (WT) controls, as indicated by lateral root number, root length, and root surface area (Fig. 2A–C).

**Figure 2.**
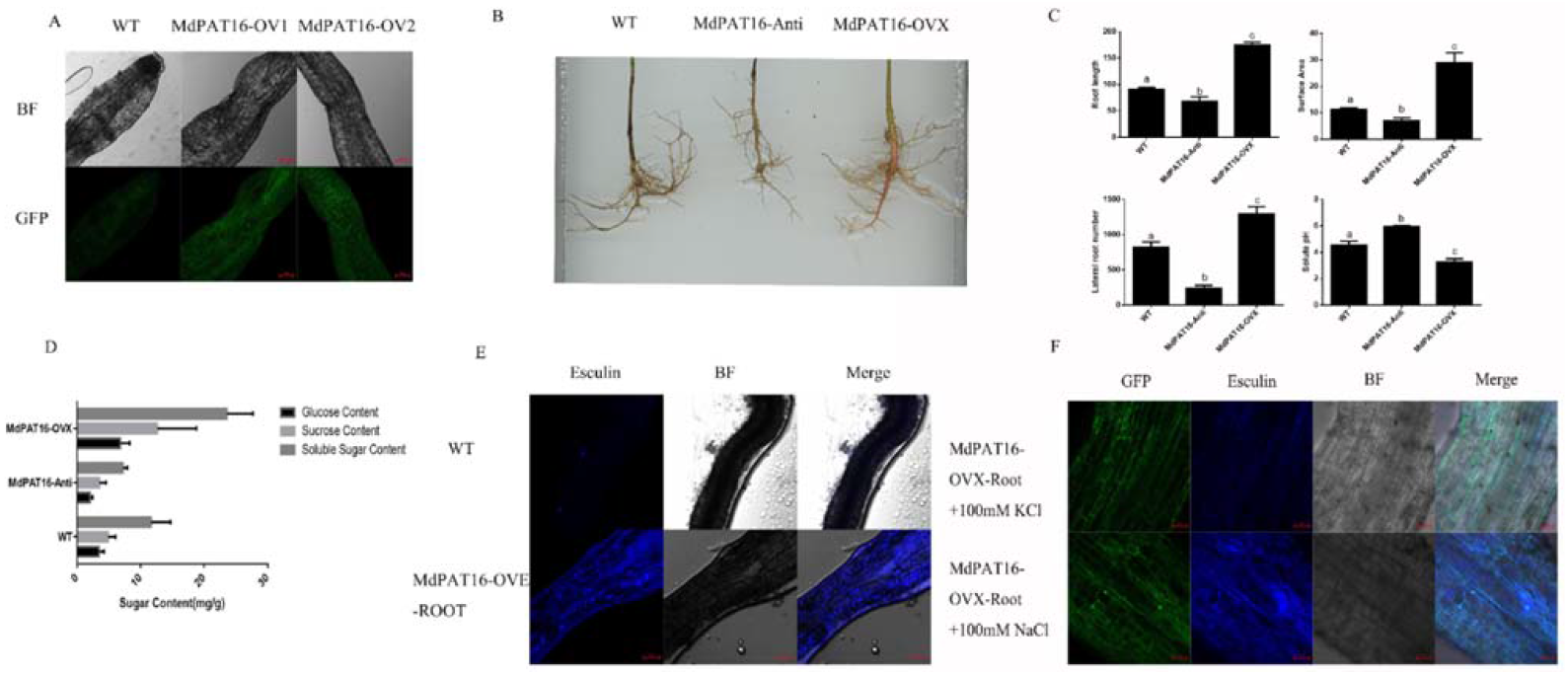
MdPAT16 functions as a positive regulator under salt stress. **(A)** Root fluorescence identification of MdPAT16 overexpression transgenic roots (Bars = 100 μm). **(B) and (C)** Root scan analysis of root shape, root length, surface area, and lateral root number of WT, MdPAT16-OVX, and MdPAT16-anti transgenic roots. Results are given as mean+SD. Letters indicate significant differences (*t*-test, *P* < 0.01) **(D)** Glucose, sucrose, and soluble sugar contents of WT, MdPAT16-OVX, and MdPAT16-anti transgenic roots. **(E) and (F)** Esculin staining of sucrose transport activity in (E) WT and MdPAT16-OVX transgenic roots, and (F) MdPAT16-OVX roots treated with NaCl and KCl (Bars = 100 μm).

Esculin staining was performed to examine sucrose transport activity in the different transgenic roots. MdPAT16-OVX roots exhibited much stronger blue fluorescence than the WT controls, indicating that MdPAT16 overexpression promoted sucrose transport activity. NaCl treatment further enhanced the fluorescence intensity (Fig. 2 E,F). Sugar contents were also measured, and MdPAT16-OVX roots accumulated much more glucose, sucrose, and soluble sugars than the WT controls, whereas MdPAT-Anti accumulated less (Fig. 2D).

To further examine whether MdPAT16 improves sugar accumulation, TRV viral vectors were used for transient overexpression and suppression (Yuval et al., 2007). MdPAT16-TRV was transiently transformed into Gala shoot cultures to inhibit the expression of MdPAT16. The resultant transgenic shoot cultures were used to detect starch content through iodine staining. MdPAT16 silencing increased starch accumulation but decreased soluble sugar contents (Fig. 3A,B). Subsequently, a VIGS experiment was conducted using apple fruits, and the results were the same as those obtained in apple roots and seedlings (Fig. 3C,D). These results suggested that MdPAT16 responded to salt stress, improved plant salt resistance, and increased sugar content.

**Figure 3.**
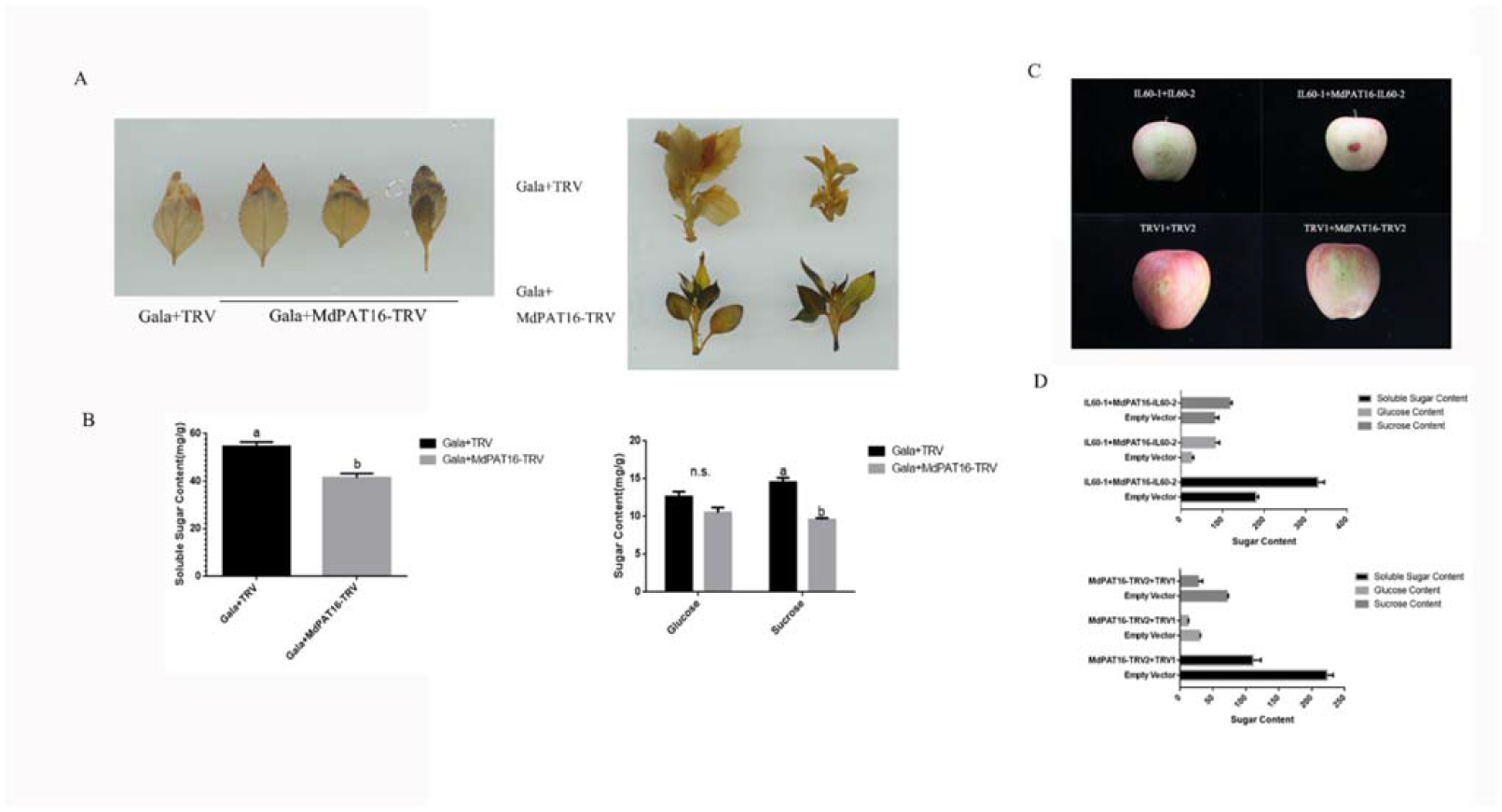
Overexpression of apple MdPAT16 increases soluble sugar content. **(A)** Starch staining of MdPAT16-TRV transient suppression Gala tissue culture seedlings and Empty Vector controls. **(B)** Soluble sugar, glucose, and sucrose contents of MdPAT16-TRV and Empty Vector controls. **(C) and (D)** visible anthocyanin accumulation conditions(C) and soluble sugar, glucose, and sucrose contents (D) of apple fruits from MdPAT16-IL60 (overexpression), MdPAT16-TRV (suppression) and Empty Vector controls.

To further characterize its function *in planta*, MdPAT16 was genetically transformed into Arabidopsis. Esculin staining demonstrated that ectopic expression of MdPAT16 enhanced fluorescence intensity compared with the WT control. As a result, ectopic transgenic lines generated more sugars than the WT controls. MdPAT16 ectopic expression also promoted salt tolerance in transgenic *Arabidopsis* (Fig. S3).

Taken together, these results indicate that *MdPAT16* plays a crucial role in sugar accumulation in response to salt stress and positively regulates salt tolerance.

### MdPAT16 is an S-palmitoyltransferase

A phylogenetic tree and a sequence alignment analysis demonstrated that *MdPAT16* is a member of the PAT gene family and that PAT16 sequences are highly conserved among different plant species (Figs. S4,S5). The yeast mutant akr1p, which is deficient in palmitoyltransferase activity, was used to determine whether MdPAT16 has palmitoylation activity. An MdPAT16-pYES2 expression vector was constructed and genetically transformed into the akr1p mutant and the WT strain BY4741, and the empty vector was used as a control. MdPAT16 ectopic expression in the akr1p mutant recovered its palmitoyltransferase deficiency phenotype, whereas expression of the empty vector did not. However, MdPAT16 ectopic expression in the akr1p mutant failed to recover its temperature sensitive phenotype.

A mutation from cysteine (Cys) to alanine (Ala) was created in the DHHC-CRD domain of MdPAT16 to produce MdPAT16^C244A^, and MdPAT16^C244A^ lost the ability to complement the palmitoyltransferase deficiency phenotype (Fig. 4A).

**Figure 4.**
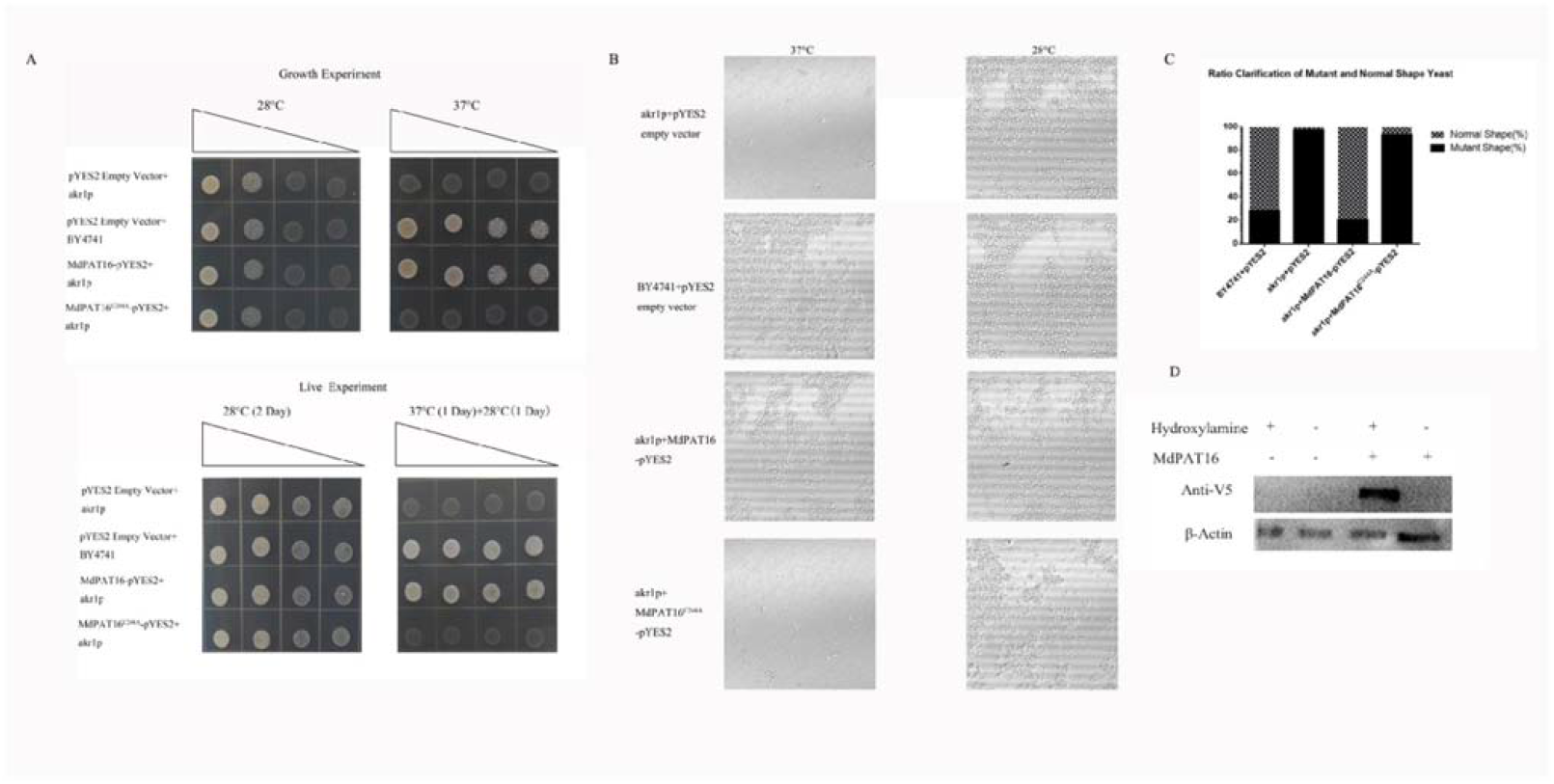
MdPAT16 is an S-palmitoyltransferase. **(A)** Growth test (above) and survival test (below). Under the nonpermissive temperature of 37°C (right), wild-type (BY4741) yeast cells grew well, but akr1p did not. Expression of MdPAT16 in akr1p largely restored growth, but MdPAT16^C244A^ failed. **(B) and (C)** Observations of cell shape using a LSM880 high resolution laser confocal microscope (B). The cell shape of MdPAT16/akr1p was indistinguishable from WT, whereas that of MdPAT16^C244A^ differed. Quantitative statistics of all four genotypes grown at 37°C (C). **(D)** MdPAT16-V5/akr1p and pYES2-V5/akr1p by ABE assay were measured by Western blotting with anti-V5 antibody to demonstrate that apple MdPAT16 is auto-acylated.

The shapes of yeast cells under different treatments were observed with an LSM880 high resolution laser confocal microscope, and quantitative statistics were used to analyze the the percentage of cells that had an altered shape. Observations showed that the expression of MdPAT16 in *akr1p* yeast cells produced a full oval shape similar to that of WT BY4741, whereas *akr1p* yeast cells that expressed MdPAT16^C244A^ exhibited a long rod shape. These observations suggested that MdPAT16 rescued the phenotype of *akr1p* and that this rescue required the cysteine residue of the DHHC catalytic site (Fig. 4B,C). Finally, an ABE (Acyl-Biotin Exchange) assay demonstrated that MdPAT16 had the capacity to palmitoylate itself (Fig. 4D). These observations strongly suggest that MdPAT16 is an S-palmitoyltransferase.

### MdPAT16 interacts with the calcineurin B subunit protein MdCBL1

To identify MdPAT16-interacting proteins *in planta*, total protein was extracted from 35S::MdPAT16-GFP and 35S::GFP transgenic apple calli for co-immunoprecipitation (Co-IP) assays. The IPed proteins were analyzed with mass spectrometry, and several identified peptides were parts of a calcineurin B subunit protein homologous to CBL1, hereafter named MdCBL1. Subsequently, 35S::MdPAT16-GFP/MdCBL1-HA and MdPAT16-GFP/HA double transgenic apple calli were obtained to verify the proteins’ interaction *in vivo* using Co-IP. The results showed that MdPAT16 interacted with MdCBL1 (Fig. 5A). Furthermore, both *in vitro* pull-down assays and bimolecular fluorescence complementarity (BiFC) assays confirmed the *in vitro* and *in vivo* interaction between MdCBL1 and MdPAT16 (Fig. 5B,C).

**Figure 5.**
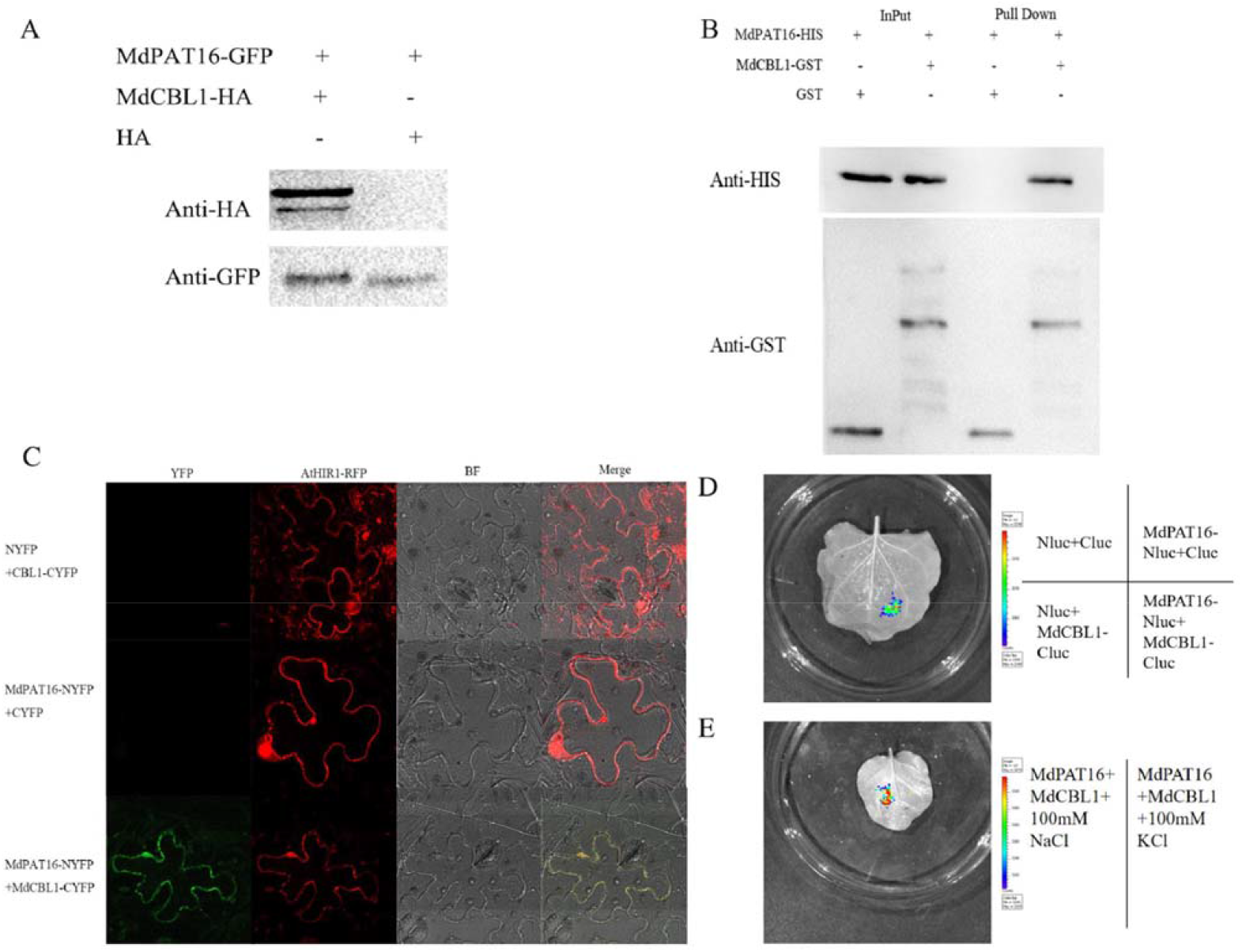
MdCBL1 is a direct substrate of MdPAT16. **(A)** *in vivo* Co-IP assays between MdPAT16 and MdCBL1 by western blotting with anti-HA and anti-GFP antibodies. **(B)** *in vitro* GST pull-down assays with MdPAT16-HIS and MdCBL1-GST. Proteins immunoprecipitated with GST-beads were detected using anti-HIS antibody. **(C)** BiFC was performed to test the interaction between MdPAT16 and MdCBL1, and AtHIR1-RFP was co-injected as a plasma membrane marker. Bars = 10 μm. **(D) and (E)** The interaction between MdPAT16 and MdCBL1 was visualized with a dual-luciferase reporter system (D) and fluorescence activity was observed with and without NaCl (E).

Next, a dual-luciferase reporter system was used to determine whether NaCl treatment influenced the interaction between MdPAT16 and MdCBL1. Full-length cDNAs of MdPAT16 and MdCBL1 were fused to the N- and C-terminals of the binary fluorescent vector, respectively. Fluorescence observations demonstrated that the luciferase activity around the MdPAT16-Nluc and MdCBL1-Cluc co-injection site was much higher than that observed in the empty vector control, further confirming the interaction between MdPAT16 and MdCBL1 (Fig. 5D). Subsequently, 100mM KCl or 100mM NaCl was added to the injection solution and co-injected with MdPAT16-Nluc and MdCBL1-Cluc. Compared with the KCl treatment, the NaCl treatment clearly promoted the interaction between MdPAT16 and MdCBL1 (Fig. 5E).

### MdPAT16 palmitoylates MdCBL1 to determine its subcellular localization and protein stability

To determine whether MdPAT16 palmitoylates MdCBL1, MdPAT16-OVX/MdCBL1-HA and MdPAT16-RNAi/MdCBL1-HA double transgenic apple calli were obtained and used for Co-IP assays with an anti-HA antibody. MdCBL1 palmitoylation was undetectable in MdPAT16-RNAi transgenic apple calli, whereas MdCBL1 was markedly palmitoylated in MdPAT16-overexpressing apple calli (Fig. 6A), indicating that MdPAT16 is required for MdCBL1 palmitoylation. Furthermore, an online palmitoylation site prediction website, CSS-Palm (http://csspalm.biocuckoo.org/), was used to predict the possible palmitoylation sites of MdCBL1. Two Cys residues, the 3^rd^ and 138^th^ cysteines, were scored as possible palmitoylation sites. Functional complementation assays of the yeast *akr1p* mutant demonstrated that MdCBL1 protein that contained mutations at each possible palmitoylation site failed to rescue the deficient phenotype (Fig. 6B), indicating that the two palmitoylation sites are crucial for MdCBL1 function.

**Figure 6.**
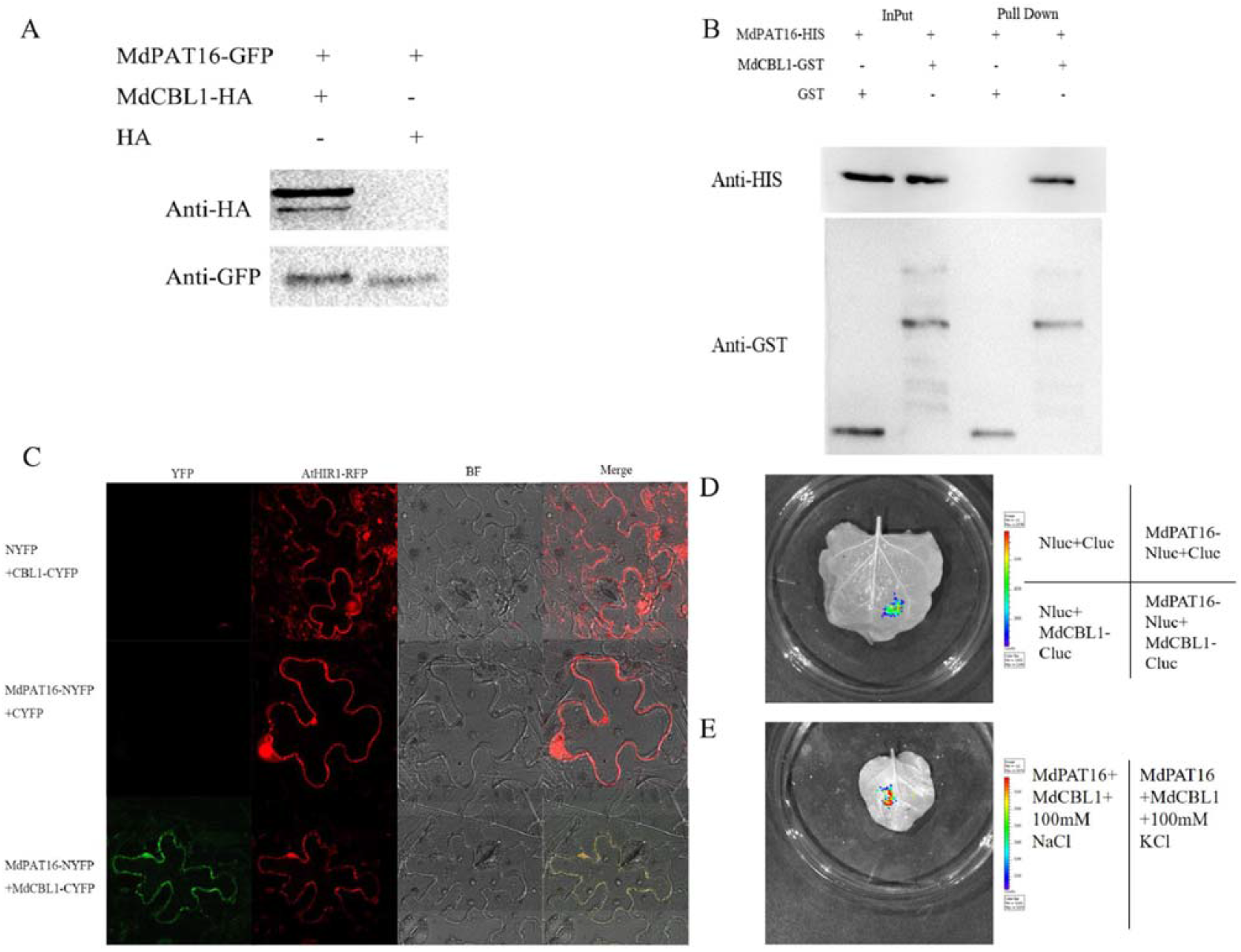
MdCBL1 is palmitoylated by MdPAT16 on the 3^rd^ cysteine residue. **(A)** *in vivo* Co-IP assays between MdPAT16/MdCBL1-HA and MdPAT16-RNAi/MdCBL1-HA double transgenic apple calli. The palmitoylation of MdCBL1 was detected using anti-HA antibodies. **(B)** Yeast functional complementation assays using MdCBL1^C3S^/akr1p and MdCBL1^C138S^/akr1p demonstrated that neither point mutant of MdCBL1 was auto-acylated. **(C)** ABE assays demonstrated that the 3rd but not 138th cysteine residue determined the palmitoylation of MdCBL1.

Acyl-biotin exchange (ABE) assays were performed to further verify the palmitoylation sites. Western blotting indicated that MdCBL1^C3S^, but not MdCBL1^C138S^, failed to be palmitoylated by MdPAT16, indicating that the 3^rd^ cysteine residue, but not the 138^th^ cysteine residue, was the palmitoylation site for the MdCBL1 protein (Fig. 6C).

To determine whether MdPAT16-mediated palmitoylation of MdCBL1 influenced its subcellular localization, MdPAT16-GFP and MdCBL1-RFP were transiently expressed in *Nicotiana benthamiana* leaves. AtCBL1-GFP and AtHIR1-RFP fluorescent proteins were used as plasma membrane markers to co-localize with MdPAT16 and MdCBL1. Both MdPAT16 and MdCBL1 showed a significant co-localization with the plasma membrane marker (Fig. 7A). MdCBL1^C3S^-RFP was then used to check whether MdPAT16-mediated MdCBL1 palmitoylation influenced its subcellular localization. MdCBL1-RFP was localized to the plasma membrane in the presence of MdPAT16-GFP, whereas MdCBL1^C3S^-RFP was not (Fig. 7B). In addition, an uptake assay was performed to further confirm the subcellular migration of MdCBL1. Cellular compartments such as nuclei, cytoplasm, and membrane structures were isolated from tobacco leaves that transiently expressed MdCBL1-GFP and MdCBL1^C3S^-GFP. Western blotting assays were performed to measure the expression levels of MdCBL1-GFP and MdCBL1^C3S^-GFP in nuclei, cytoplasm, and membrane structures. Anti-Histone3(H3) and anti-actin were used as loading controls for nuclei and cytoplasm, respectively, while AtHIR1-RFP was co-injected as a loading control for membrane structures. The results indicated that the subcellular localization of MdCBL1 changes when palmitoylation is absent (Fig. 7C).

**Figure 7.**
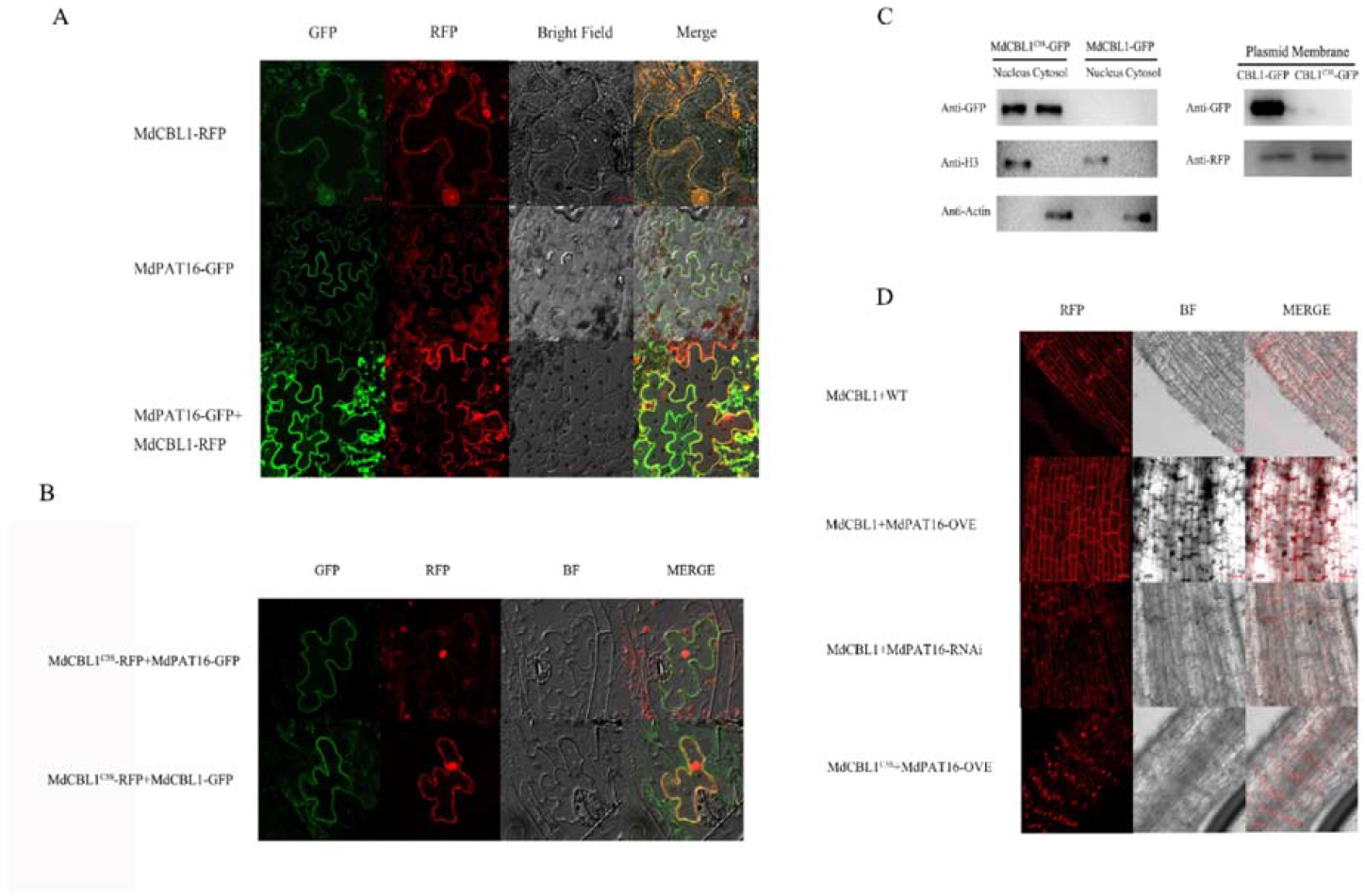
Localization of MdCBL1 to the plasma membrane depends on the function of MdPAT16. **(A)** Subcellular localization of MdCBL1-RFP and MdPAT16-GFP in *N. benthamiana* leaves. AtCBL1-GFP and AtHIR1-RFP were used as plasma membrane markers. **(B)** Subcellular localization of MdCBL1^C3S^-RFP. **(C)** Qualitative detection of MdCBL1 and MdCBL1^C3S^ in different cellular compartments by western blotting. Anti-Histone3 and anti-actin were used as loading controls for nuclei and cytoplasm, respectively. AtHIR1-RFP was co-injected as a loading control for plasma membrane. **(D)** Subcellular localization of MdCBL1/WT, MdCBL1/MdPAT16-OVE, MdCBL1/MdPAT16-RNAi, and MdCBL1^C3S^/MdPAT16-OVE in transgenic roots of GL-3.

**Figure 8.**
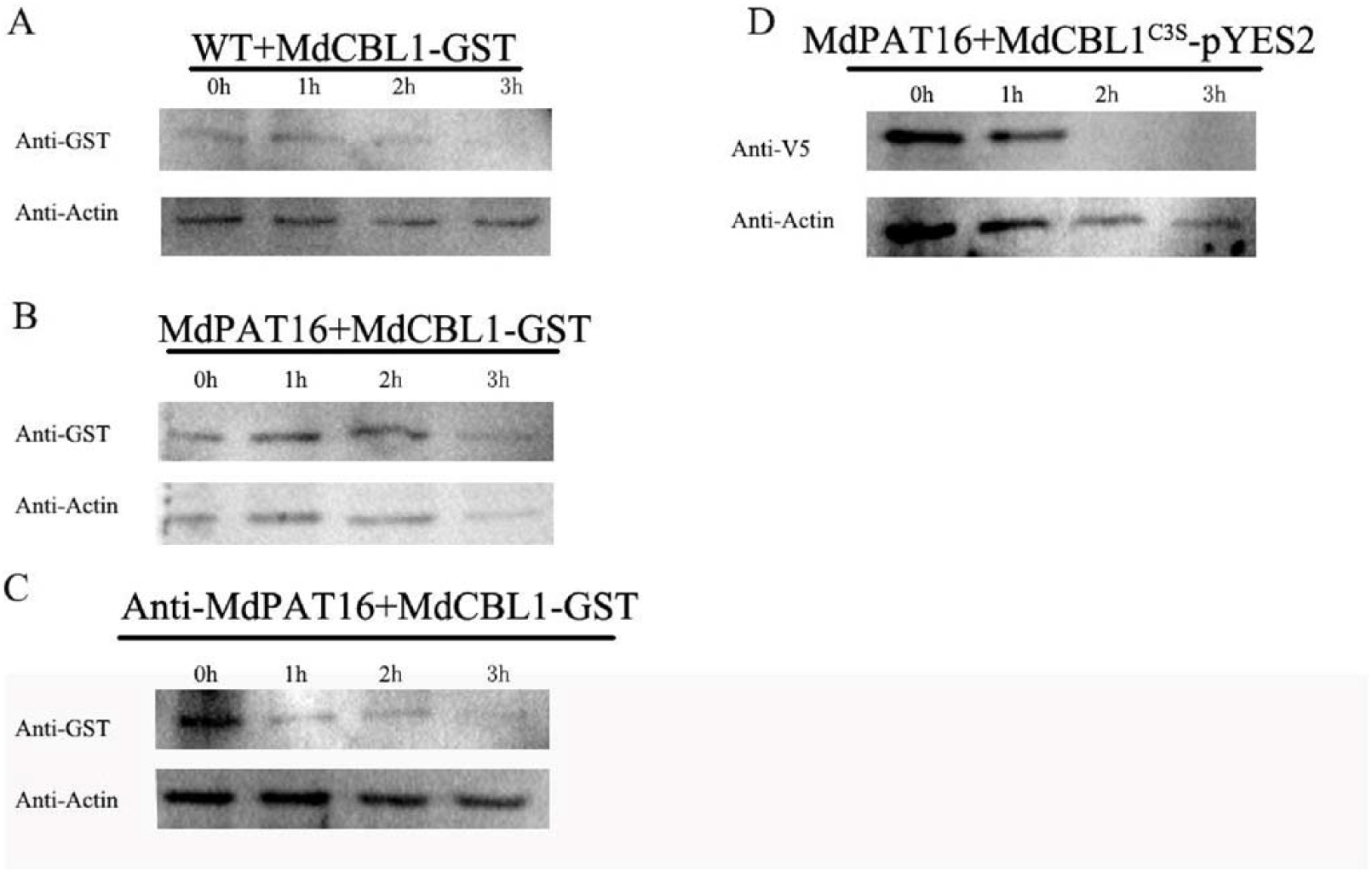
MdPAT16 stabilizes MdCBL1 through palmitoylation. **(A), (B) and (C)** Cell-free degradation assay in which MdCBL1-GST was recombined with total proteins extracted from WT (A), MdPAT16-OVE (B), and MdPAT16-RNAi (C) apple calli under a water bath for 1, 2, and 3 h. The expression of MdCBL1 was measured by western blotting. **(D)** Cell-free degradation assay in which MdCBL1^C3S^-V5 was recombined with total proteins extracted from MdPAT16-OVE apple calli under a water bath for 1, 2, and 3 h.

In addition, 35S::MdPAT16-OVE/MdCBL1-RFP, 35S::MdPAT16-RNAi/MdCBL1-RFP, 35S::Empty Vector/MdCBL1-RFP, and 35S::MdPAT16-OVE/MdCBL1^C3S^-RFP were genetically transformed into apple roots using an *Agrobacterium rhizogene*s-mediated transformation system. The resultant transgenic roots were used to determine whether the MdPAT16-mediated palmitoylation of MdCBL1 influences its subcellular localization. MdPAT16 was required for the plasma membrane localization of MdCBL1-RFP protein, and MdCBL1^C3S^-RFP failed to localize to plasma membrane even together with MdPAT16 in 35S::MdPAT16-OVE/MdCBL1^C3S^-RFP transgenic apple roots (Fig. 7D). Therefore, MdPAT16-mediated palmitoylation of MdCBL1 promotes its subcellular localization to the plasma membrane.

Cell-free degradation assays were performed to examine whether MdPAT16-mediated palmitoylation affects the protein stability of MdCBL1. Prokaryotic-expressed MdCBL1-GST and eukaryotic-expressed MdCBL1^C3S^-V5 were incubated with total proteins extracted from the WT control and from MdPAT16-GFP and MdPAT16-RNAi transgenic apple calli. MdCBL1-GST was more stable in MdPAT16-GFP transgenic apple calli, but its stability was markedly decreased in MdPAT16 RNAi transgenic calli. Meanwhile, MdCBL1^C3S^-V5 degraded much more quickly than MdCBL1-GST. Therefore, MdPAT16-mediated palmitoylation of MdCBL1 influences its stability.

### MdCIPK13 is required for MdPAT16-mediated sugar accumulation

To determine how MdPAT16 and MdCBL1 are involved in the regulation of sugar accumulation, viral vectors were used to perform transient expression analyses in apple fruits. MdCBL1-IL60 overexpression and MdCBL1-TRV suppression vectors were constructed and injected into fruits. Like MdPAT16, MdCBL1 overexpression clearly increased sugar content, whereas its suppression decreased sugar content. As mentioned above, MdPAT16 overexpression increased sugar content (Figure 3C,D). When MdPAT16-IL60 and MdCBL1-TRV were co-injected into apple fruit, MdCBL1 suppression almost completely abolished the MdPAT16-mediated increase in fruit sugar content (Fig. 9A-C). In addition, *Agrobacterium rhizogene*s-mediated genetic transformation was performed to obtain 35S::MdCBL1-OVX, 35S::MdCBL1-RNAi, and 35S::MdCBL1^C3S^-OVX transgenic apple roots. Increasing activity of sucrose transferase was detected in the resultant transgenic roots, respectively. MdCBL1 overexpression enhanced sucrose transferase activity in transgenic apple roots, and MdCBL1 suppression inhibited it (Fig. 9D). Furthermore, mutation of the MdCBL1 palmitoylation site abolished its role in enhancing sucrose transferase activity (Fig. 9D), indicating that the palmitoylation site plays a crucial role in MdCBL1 function.

**Figure 9.**
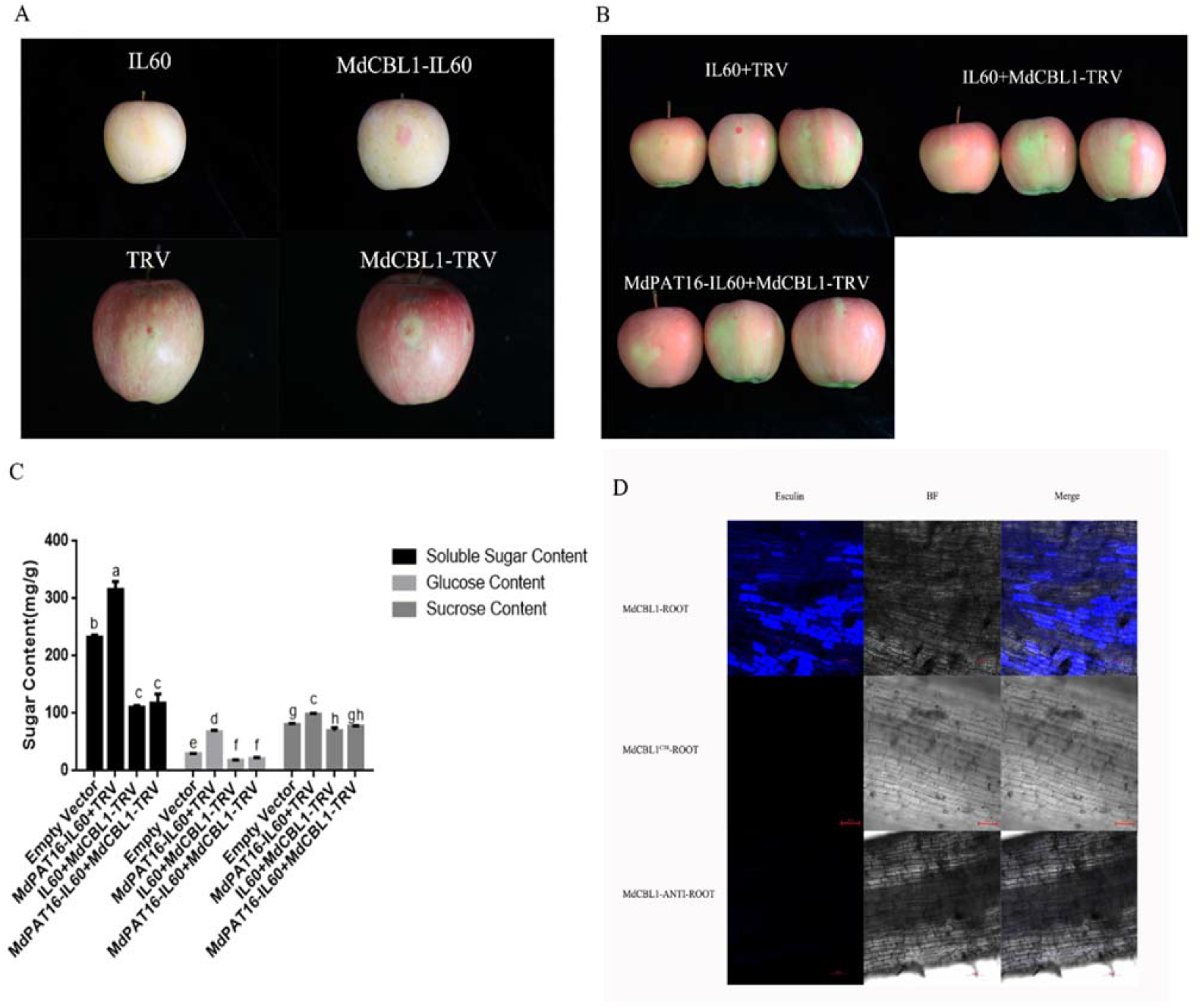
MdCBL1 is required for sugar accumulation. **(A) and (B)** Anthocyanin accumulation of MdCBL1-IL60, MdCBL1-TRV and Empty Vector apple fruits (A), and MdPAT16-IL60/MdCBL1-TRV, IL60/MdCBL1-TRV, and Empty Vector apple fruits (B). **(C)** Soluble sugar, glucose, and sucrose contents of different fruits. **(D)** Esculin uptake assay for sucrose transport activity in transgenic roots of MdCBL1, MdCBL1^C3S^, and MdCBL1-RNAi.

In our previous report, salt stress induced sugar accumulation in the vacuole through a MdCBL1/MdCIPK13-MdSUT2.2 pathway (Ma et al., 2019). Considering that MdCBL1 interacts with MdCIPK13, it is reasonable to propose that MdCIPK13 is involved in MdPAT16-mediated sugar accumulation in response to salt stress. To verify this hypothesis, viral vectors were used to perform transient expression experiments in apple fruits. MdCIPK13-IL60 overexpression and MdCIPK13-TRV suppression vectors were constructed and used for injection. MdCIPK13-IL60 promoted sugar accumulation in apple fruit, whereas MdCIPK13-TRV inhibited it (Fig. 10A). In addition, MdCIPK13 suppression abolished the MdPAT16-mediated sugar increase in MdPAT16-IL60+MdCIPK13-TRV co-injected apple fruits (Fig. 10A,B), indicating that MdCIPK13 is required for MdPAT16-mediated sugar accumulation under salt stress.

**Figure 10.**
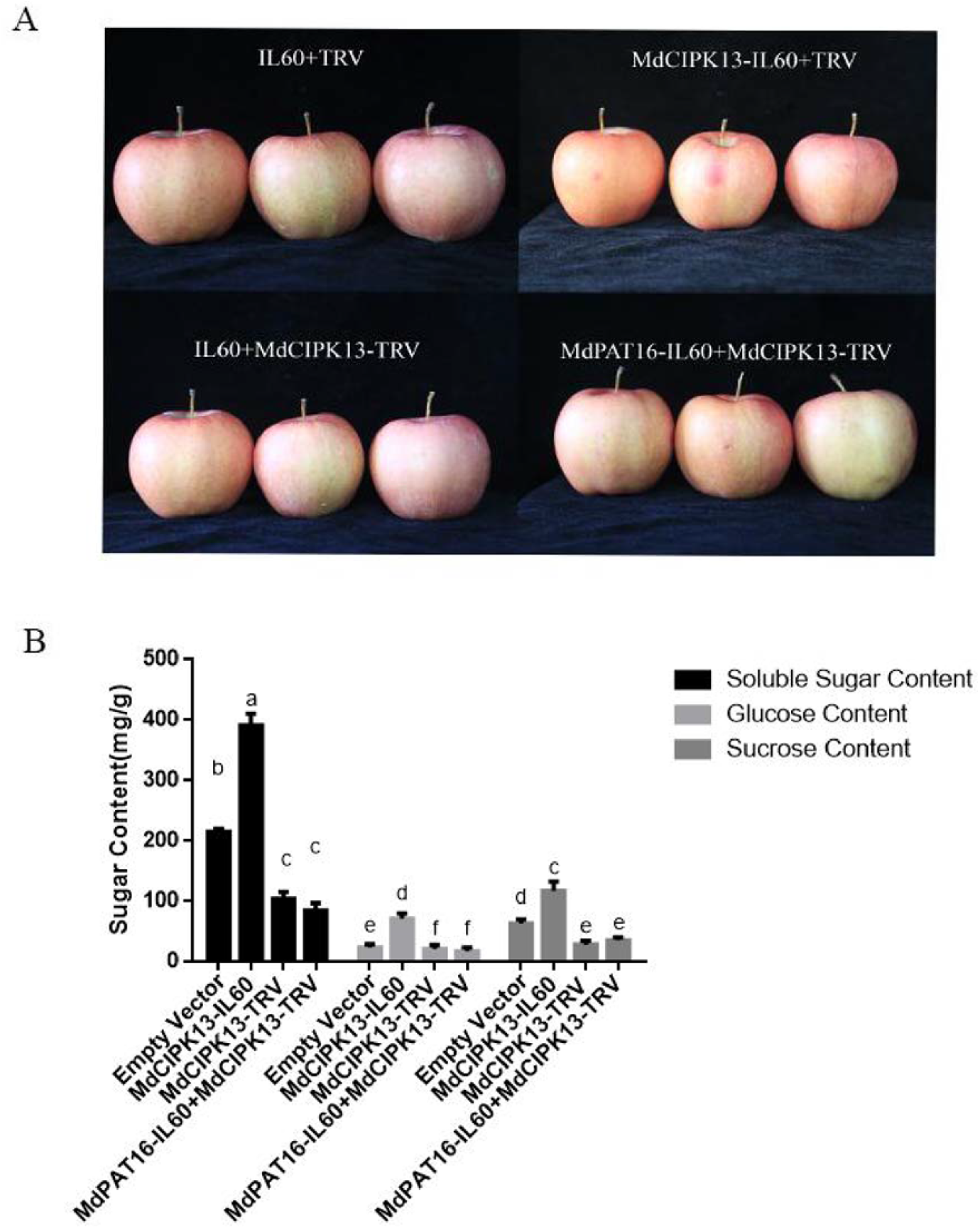
MdPAT16 mediates sugar accumulation through the MdCIPK13 pathway. Anthocyanin accumulation **(A)** and soluble sugar, glucose, and sucrose contents **(B)** of MdCIPK13-IL60, MdCIPK13-TRV, MdPAT16-IL60/MdCIPK13-TRV, and Empty Vector apple fruits.

## Discussion

Salt stress impedes plant growth and development via osmotic adjustment (Munns, 2002). It also alters stomatal conductance and transpiration rate and affects plant ion balance (Chaves et al., 2009). Soluble sugars, as compatible osmolytes, are thought to function mainly in the stabilization of proteins and cellular structures under stress (Sanchez et al., 2008). In addition, sugars reduce the stress-induced accumulation of reactive oxygen species (ROS) (Hu et al., 2012), thereby increasing stress resistance. Although attention has been paid to the function of sugars under salt stress, the specific mechanisms and upstream regulation of this process are still unclear. In this study, a palmitoylation transferase family member, MdPAT16, was identified in apple. In transgenic materials and VIGS fruits, MdPAT16 overexpression significantly increased soluble sugar content and root sucrose transferase activity under salt stress (Fig. 3), thereby positively modulating sugar accumulation and salt tolerance. A calcineurin B-like (CBL) protein, MdCBL1, was shown to be a substrate of MdPAT16. Further experiments demonstrated that MdPAT16 participated in the MdCBL1-MdCIPK13-MdSUT2.2 regulatory pathway through palmitoylation of MdCBL1, thereby regulating sugar accumulation (Figs. 9,10; Ma et al., 2019a).

PATs respond to abiotic stress. The loss-of-function mutant *pat10* exhibits hypersensitivity to salt stress in *Arabidopsis thaliana* (Cheng et al., 2005; Krebs et al., 2010; Bassil et al., 2011b; Barragán et al., 2012; Zhou et al., 2013). In maize, the S-palmitoylation family member ZmTIP1 interacts with the calcium-dependent protein kinase ZmCPK9 to regulate root hair length and drought tolerance (Zhang et al., 2019). In apple, MdPAT16 was identified as a DHHC-type PAT (Figure S5). It played roles not only in the modulation of salt tolerance (Figure 2), but also in the regulation of sugar accumulation (Figure 3).

The critical functions of PATs do not depend exclusively on the DHHC functional group, and not all DHHC-type PATs show palmitoyltransferase activity. Some PAT activity deficient proteins also function to the resistance to external stress, and PATs can act as cation transporters to withstand adverse circumstances (Hines et al., 2010; Ohno et al., 2012). Indeed, we could not initially exclude the possibility that the functions of MdPAT16 in salt stress resistance and sugar accumulation derived not from its palmitoylation of substrates but from its potential role as a cation transporter. However, functional complementation assays demonstrated that MdPAT16 rescued the palmitoyltransferase deficient phenotype of the akr1p yeast mutant, whereas MdPAT16^C244A^ failed, indicating that MdPAT16 is a typical palmitoyltransferase. These results suggested that MdPAT16 functions through its PAT activity.

The functions of PATs are usually achieved through palmitoylation of substrate proteins. Palmitoylation regulates protein activity, protein sorting, and protein-protein interactions (Hemsley et al., 2013; Zhou et al., 2013). Palmitoylation also influences protein stabilization, promoting the function of substrate proteins. The human palmitoyltransferase ZDHHC14 has been shown to exert a tumor suppressor function through palmitoylation of its substrate proteins (Marc et al., 2014). A series of calmodulin proteins in plants that respond to calcium signals have been identified as PAT substrates, including CPKs (Leclercq et al., 2005; Martín and Busconi, 2000; Zhang et al., 2019), CaMs (Wang et al., 2005), and CBLs (Batistič et al., 2010; Zhou et al., 2013). Here. the CBL family protein MdCBL1 was identified as a direct substrate of MdPAT16 using mass spectrometry. MdCBL1 directly interacted with MdPAT16 *in vivo* and *in vitro*. Experiments demonstrated that the palmitoylation of MdCBL1 was dependent on MdPAT16, and MdCBL1 created a phenotype indistinguishable from that of MdPAT16 (Fig. 5). Therefore, MdCBL1 was a direct substrate of MdPAT16.

Substrate proteins lose membrane localization and biological function in the absence of corresponding PAT modifications. CBL2, 3, and 6 in *Arabidopsis thaliana* are substrates of PAT10, and palmitoylation site deletion mutants of CBL2, 3, and 6 fail to localize to the tonoplast (Batistič, 2012; Zhou et al., 2013). The *cbl2* and *cbl3* double mutant exhibits developmental abnormalities, including leaf tip necrosis and defects in the reproductive process, that are similar to the phenotype of the *pat10* single mutant (Tang et al., 2012; Zhou et al., 2013). In apple, the membrane localization and function of MdCBL1 also depended on its palmitoylation. The palmitoylation site mutant MdCBL1^C3S^ no longer localized to the plasma membrane, and 35S-driven MdCBL1^C3S^ transgenic roots showed a functional absence of sucrose transferase activity without corresponding PAT modifications (Figs. 7,9D). Thus, palmitoylation modification played an important role in the function of substrate proteins. Taken together, these results suggest that the function of MdCBL1 in the regulation of sugar content is dependent on palmitoylation by MdPAT16.

Transgenic analyses suggested that MdCBL1^C3S^ was mislocalized from the plasma membrane to the cytoplasm and nucleus (Fig. 7) and lost its regulatory function in sugar accumulation. Ubiquitination in plants modulates the nuclear entry and degradation of substrate proteins. Therefore, a ubiquitination enzyme may facilitate the nuclear entry and degradation of MdCBL1. Studies in humans indicate that a competitive inhibition exists between palmitoylation and ubiquitination of substrate proteins, in which palmitoylation inhibits the activity of ubiquitination enzymes (Rebecca et al., 2019). The palmitoylation modifications of PD-L1 by DHHC3 significantly inhibit the ubiquitination modifications of PD-L1 (Han et al., 2019). Therefore, MdCBL1 may be modified by ubiquitination in the absence of appropriate palmitoylation. This ubiquitination modification may be directly or indirectly affected by palmitoylation.

The biological functions of CBLs are performed through interaction with CIPKs (Cheong, 2003; Li et al., 2006; Pandey et al., 2004). In Arabidopsis, AtCBL1 shows a conserved interaction with AtCIPK24/SOS2 that mediates sodium ion homeostasis under salt stress (Albrecht et al., 2003). In *M. domestica*, the overexpression of MdCIPK6L causes significant improvements in salt tolerance, and the heterologous expression of MdCIPK6L rescues the salt-sensitive phenotype of *sos2* (Wang et al., 2012). MdSOS2 exhibits high similarity with AtCIPK24/SOS2, which also shows high salt tolerance (Hu et al., 2012). Apple MdSUT2.2 enhances sugar accumulation and stress resistance by the phosphorylation of MdCIPK13 (Ma et al., 2019a). Our results suggest that MdPAT16 is involved in the MdCIPK13 regulatory pathway through its interactions with MdCBL1. The palmitoylation stabilization of MdCBL1 probably stabilizes the MdCBL1-MdCIPK13 protein-protein interaction and promotes the phosphorylation of MdSUT2.2, thereby causing sugar accumulation. Previous studies have also demonstrated that another *M. domestica* CIPK family member, MdCIPK22, functions similarly to MdCIPK13, interacting with MdSUT2.2 in response to drought stress to promote sugar accumulation (Ma et al., 2019b). Whether PAT is also involved in the upstream regulation of this process requires further investigation.

MdCIPK13 and MdSUT2.2 are co-localized on the tonoplast, unlike MdCBL1 and MdPAT16 that co-localize to the plasma membrane. Studies on AtCBL1 demonstrate that when AtCIPK1 interacts with plasma-membrane-localized AtCBL1 or tonoplast-localized AtCBL2, the subcellular localization of AtCIPK1 changes correspondently, resulting in different membrane localization (Batistič et al., 2008). A similar situation may occur for MdCIPK13, which exhibits tonoplast localization when interacting with MdSUT2.2 and plasma membrane localization when interacting with MdCBL1. Therefore, MdCIPK13 probably migrates during the interaction with different proteins, resulting in different localizations. Further experiments should be performed to assess this hypothesis.

Taken together, previous reports and our current study suggest a working model that summarizes our findings. Under salt stress in apple, MdPAT16 stabilizes the expression of MdCBL1 through palmitoylation and promotes sugar accumulation through the regulatory pathway of MdCBL1-MdCIPK13-MdSUT2.2 (Fig. S6). Meanwhile, there exists a ubiquitin ligase that mediates the nuclear entry and degradation of MdCBL1 in the absence of palmitoylation. This process may be directly or indirectly regulated by palmitoylation.

Sugar content in apple is a decisive factor for fruit quality. Sugars accumulate to maintain the intracellular ion balance in response to salt stress and also contribute to fruit quality and commodity values (Hu et al., 2016). Our work reveals a new mechanism for the regulation of sugar content and fruit quality and contributes to a better understanding the pathway by which apple responds to salt stress. Increases in carbohydrate concentration following moderate salt stress raise the question of the role of carbohydrate availability in plant growth under stress. In total, our data support the proposal that moderate upregulation of MdPAT16 has a potential role in the promotion of fruit quality during salt stress. Therefore, our research provides new ideas for the simultaneous improvement of fruit quality and stress resistance through breeding methods.

## Materials and methods

### Plant material, growth conditions, and salt treatments

GL-3 apple (*Malus x domestica* Borkh.) tissue cultures were used for transformation and stress treatments (Chen et al., 2019). GL-3 and transgenic apple cultures were cultured and subcultured in Murashige & Skoog (MS) medium supplemented with 0.3 mg/L 6-benzylaminopurine (6-BA), 0.2 mg/L indoleacetic acid (IAA), and 0.1 mg/L gibberellin (GA). The cultures were maintained at a constant temperature of 25°C under long-day conditions (16 h light/8 h dark) and were subcultured every 30 days.

Seeds of *Malus hupehensis* were harvested and stratified at low temperature and humidity for more than 30 days. When the seeds began to germinate, they were transplanted into substrate under long-day greenhouse conditions (16 h light/8 h dark). Four weeks after germination, seedlings of consistent size were selected for further experiments.

Apple calli used in this article were induced from embryos of ‘Orin’ apples (*Malus x domestica* Borkh.). Calli were grown and subcultured on MS medium with 1.5 mg/L 2,4-dichlorophenoxyacetic acid and 0.5 mg/L 6-BA at a constant temperature of 25°C in the dark and subcultured every 15 days.

For short-term salt treatments, 150 mM NaCl was used in hydroponic experiments. For long-term salt treatments, apple plantlets were treated with 150 mM NaCl under the greenhouse conditions described above.

Different concentrations of NaCl(0mM; 1mM; 10mM; and 100mM) were applied to *Malus hupehensis* apple seedlings after germination and transplanting into vermiculite. The pictures were obtained after two weeks under the same greenhouse conditions described above.

### Ca^2+^-ATPase activity, Rhizosphere pH staining, and Solution pH

The activities of Ca^2+^-ATPase in *Malus hupehensis* apple seedlings were detected by colorimetry method. The method was followed by Ca^2+^/Mg^2+^-ATPase activity detection kit(BC0965, Solarbio).

The treated roots of *Malus hupehensis* apple seedlings were placed in the solution mixed of 0.01% bromocresol violet, 0.2mm CaSO_4_, and 0.7% Agar (pH = 6.5), and photographes were taken after 45 minutes in dark.

Solution pH was detected by PHS-3C pH-meter.

### Genetic transformation

The full-length coding sequences (CDS) of *MdPAT16* (MD10G1058600) and *MdCBL1* (MD00G1132600) were identified from the apple genome (GDDH13 v1.1) and amplified. The CDS of *MdPAT16* and *MdCBL1* were inserted into the pRI101-An plasmid with a GFP tag to build overexpression vectors. The forward and reverse fragments were also inserted into the pRNAi-E vector (Song et al., 2017) to construct RNAi vectors. Both overexpression vectors and RNAi vectors were introduced into *Agrobacterium Tumaficiens* EHA105 competent cells (Transgen). The genetic transformation procedure was performed as described in Ma et al. (2017b).

The full length *MdPAT16* sequence was then ligated with a GFP tag to build MdPAT16-GFP, and the full length *MdCBL1* sequence was ligated with an HA tag to build MdCBL1-HA. These vectors were introduced into ‘Orin’ apple calli using the Agrobacterium method as described in Ma et al. (2017a).

Transgenic root systems of MdPAT16 and MdCBL1 were induced by *Agrobacterium Rhizogenesis* K599 competent cells (Transgen) under *Malus hupehensis* and GL-3 tissue culture backgrounds according to the procedure of Meng et al. (2019).

For Arabidopsis transformation, the MdPAT16 overexpression vector was introduced into *A. Tumaficiens* strain LBA4404. MdPAT16 was then transformed into Col-0 using the floral dip method, and the seeds were screened on 1/2 MS medium with 25 mg/L kanamycin. Positive seedlings were detected using qRT-PCR, and the screened T3 generation seedlings were used for further analysis.

### RNA extraction and qRT-PCR assays

Total RNA was extracted from rooted tissue cultures using the RNA Extraction Kit (Tiangen). The extracted RNA was purified using the PrimeScript First Strand cDNA Synthesis Kit (TaKaRa, Dalian, China), and the oligo DT Kit was used to generate cDNA.

A 20 μl reaction system with SYBR Green Supermix (Takara) was used for qRT-PCR analysis. The total reaction system included 10 μl SYBR Green mixture, 7 μl double distilled water, 1 μl cDNA template, and 1 μl each of upstream and downstream primers. Relative expression was quantified using the (Ct) 2^−DDCt^ method, and *MdActin* (GenBank accession number CN938024) was selected as the internal control gene. Prism software was used to generate the chart, and the significance of treatment differences was assessed by the Data processing system software using One-way ANOVA method.

### Yeast functional complementation assay

The yeast palmitoylation mutant akr1p and its original strain, BY4741, were obtained from Thermo Scientific. The pYES2-DEST52 empty vector, MdPAT16-pYES2, and MdPAT16^C244A^-pYES2 vectors were transformed into akr1p and BY4741. For the growth assay, yeast cells were grown to stationary phase on glucose minimal liquid media. 5 μl per yeast strain with an OD_600_ of approximately 0.5 was placed onto two individual galactose minimal agar medium plates. These plates were incubated at 28 °C and 37 °C, respectively. Cells were observed under a LSM880 laser scanning confocal microscope to obtain cell shape data, and the mutant rate was counted using ImageJ (ver. 1.41).

### Acyl-Biotinyl Exchange (ABE) assay

Auto-acylation of MdPAT16 was measured using an ABE assay. MdPAT16-pYES2 and MdPAT16^C244A^-pYES2 vectors were transformed into BY4741. Total proteins were then extracted and measured by western blotting using β-actin antibody and anti-V5 antibody. The detected proteins were selected for the ABE assay following the methods of Wan et al. (2007).

### Root shape, root length, surface area, and lateral root number

The shape of roots were photographed by LA-S root scanner, and root length, surface area, and lateral root number were detected by root analyse system software by LA-S root scanner.

### Glucose; Sucrose; and Soluble sugar contents detection

The sugar contents were detected of same size (about 500mg) plant samples with three individual duplications. Sucrose, glucose, and soluble sugar assay kits(Keming biotechnology co. LTD) were used to detected sugar contents in samples.

### Esculin uptake

The roots of transgenic root systems of MdPAT16 and MdCBL1 were rinsed and mounted on glass slides in 1/2 MS liquid media with 10 mM esculin under normal or 150mM saline conditions. Fluorescence was scanned using a 367 nm excitation wavelength and a 454 nm emission wavelength as described in Ma et al. (2018).

### Virus-induced gene silencing assays

Virus-induced gene silencing (VIGS) assays were performed to verify the expression patterns of MdPAT16 and MdCBL1 in apple fruit. The full length CDS of MdPAT16 and MdCBL1 were inserted into IL60-2 vectors, to construct MdPAT16-IL60-2 and MdCBL1-IL60-2, and the IL60-1 vector was used as an auxiliary plasmid. Antisense gene fragments were also used to construct MdPAT16-TRV-2 and MdCBL1-TRV-2. The TRV1 vector was used as an auxiliary plasmid. The TRV vectors were introduced into *A. Tumaficiens* strain LBA4404. The mixed vectors and *A. Tumefaciens* solutions were injected into the peels of apple fruits, and stored under 24°C with constant light. VIGS assays were performed according to the method of An et al. (2018).

### Co-IP assay

Proteins were extracted from apple calli that were transformed with MdPAT16-GFP/HA and MdPAT16-GFP/MdCBL1-HA. The target proteins were absorbed by Protein A/G agarose beads (Thermo Fisher). The absorbed proteins were measured by western blotting using anti-MYC and anti-HA antibodies.

### Pull-down assay

The full length of CDS of MdPAT16 and MdCBL1 were ligated to His and GST targets to build MdPAT16-His and MdCBL1-GST. The resulting plasmids were transformed into *Escherichia coli* BL21 (DE3; Transgene), and 10 μM isopropyl β-D-1-thiogalactopyranoside (IPTG) was used to induce for 6 hours. Proteins were then separated and purified. The MdPAT16-His target protein was co-incubated separately with GST and MdCBL1-GST. The mixtures were eluted from glutathione-agarose beads and measured with Anti-His and Anti-GST antibodies via western blotting.

### Bimolecular fluorescence complementation (BiFC) assays

Bimolecular fluorescence complementation (BiFC) assays were performed to verify the protein-protein interaction *in vivo*. The full-length CDS of MdPAT16 and MdCBL1 were inserted into separate fluorescence plasmids to build MdPAT16-pSPYNE and MdCBL1-pSPYCE. The two vectors and the empty vector control were transformed into tobacco (*Nicotiana benthamiana*) leaves. LSM880 laser scanning confocal microscopy (488–534 nm wavelength) was used to visualize green fluorescence, and AtCBL1, a known membrane-localized protein, was used as a plasma membrane marker.

### Subcellular localization

MdPAT16 was ligated with a GFP target, MdCBL1 was ligated with an RFP tag, and MdCBL1^C3S^ was ligated with an RFP tag. The resulting plasmids were transformed into Agrobacterium LBA4404. Membrane-localized GFP and RFP proteins were used as markers. All strains were transiently transformed into *N. benthamiana* leaves to observe the single localization and co-localization conditions. The proteins were extracted from leaves to perform a separation assay, and western blotting was used to verify.

*Agrobacterium rhizogenes*-induced transgenic roots were scanned using an LSM880 laser scanning confocal microscope (488–543 nm) to measure GFP fluorescence, and FM4-64 was used as membrane-associated marker.

### Cell-free degradation

The MdCBL1-GST protein was induced and mixed with total proteins extracted from MdPAT16-OVE and MdPAT16-RNAi transgenic apple calli, with WT apple calli proteins used as a control. The reaction mixes were incubated at 22 °C and periodically sampled. The samples were boiled and measured by western blotting using anti-Actin and anti-GST antibodies.

## SUPPLEMENTAL DATA

**Supplementary Figure S1. Salt stress in apple promotes sugar accumulation.**

**Supplementary Figure S2. The MdPAT16 promoter responds to salt stress.**

**Supplementary Figure S3. Ectopic expression of MdPAT16 in Arabidopsis increases sugar content and salt resistance.**

**Supplementary Figure S4. Phylogenetic tree analysis of MdPATs and AtPATs.**

**Supplementary Figure S5. Sequence analysis of MdPAT16 and PAT16 in other species.**

**Supplementary Figure S6. Proposed model of the mechanism regulating sugar content.**

**Supplemental Table S1 The primers used in this study.**

## ACKNOWLEDGEMENTS

This work was financially supported grants from National Natural Science Foundation of China (U1706202), National Key Research and Development Program (2018YFD1000200), Ministry of Agriculture of China (CARS-27) and Shandong Province (SDAIT-06-03).

